# A Modification of the Bielschowsky Silver Stain for Alzheimer Neuritic Plaques: Suppression of Artifactual Staining by Pretreatment with Oxidizing Agents

**DOI:** 10.1101/570093

**Authors:** Anthony J. Intorcia, Jessica R. Filon, Brittany Hoffman, Geidy E. Serrano, Lucia I. Sue, Thomas G. Beach

## Abstract

The Bielschowsky silver method has been the most commonly-used histological stain for demonstrating the neuritic plaques and neurofibrillary tangles that define Alzheimer’s disease (AD). The stain has been critical to providing a common measure allowing large clinicopathological correlation studies that have demonstrated statistically significant and independent contributions of both lesion types to cognitive impairment and dementia. The continuing relevance of neuritic plaques, and the Bielschowsky stain as the method of choice for their demonstration, is also indicated by its use as the US Food and Drug Administration (FDA) “standard of truth” for autopsy confirmation of the validity of PET amyloid imaging agents. Many modifications of the Bielschowsky stain have been published but it is still known to frequently generate artifactual staining that at times mimics the appearance of amyloid plaques. In this study we found that pretreatment with oxidizing agents prior to the initial incubation in silver nitrate eliminates these artifacts and reduces the commonly observed high background staining. This new method may be valuable for Alzheimer’s disease (AD) researchers and neuropathologists.

## Introduction

The Bielschowsky silver method, first published by Max Bielschowsky in 1908, has been the most commonly used histological stain, since the time of Alzheimer, for demonstrating the senile plaques and neurofibrillary tangles that define the pathology of the disease named for him. The evolution of subsequent modifications is described in a comprehensive review by Uchihara^1^. The method was adopted in 1991 by the Committee to Establish a Registry for Alzheimer’s Disease (CERAD), together with a diagrammatic template, to provide a standardized and reproducible density assessment for both neuritic plaques and neurofibrillary tangles ^2-4^. This was critical to providing a common measure allowing large clinicopathological correlation studies that demonstrated statistically significant and independent contributions of both lesion types to cognitive impairment and dementia ^5^. Most recently, an expert committee sponsored by the National Institute on Aging and Alzheimer’s Association (NIA-AA) have revised the histopathological assessment of Alzheimer’s disease (AD) by incorporating the CERAD neuritic plaque density assessment together with Braak neurofibrillary degeneration staging and Thal amyloid phases^6, 7^ and a subsequent study of staining methods at multiple laboratories has confirmed the usefulness of the Bielschowsky stain as well as its reproducibility between centers^8^. The continuing relevance of neuritic plaques, and the Bielschowsky stain as the method of choice for their demonstration, is also indicated by its use as the US Food and Drug Administration (FDA) “standard of truth” for autopsy confirmation of the validity of PET amyloid imaging agents ^9-11^. Many modifications of the Bielschowsky stain have been published but it is still known to frequently generate artifactual staining that at times mimics the appearance of senile plaques. This most often occurs when the section has few or no true plaques or neurofibrillary change. In this study we found that pretreatment with oxidizing agents prior to the initial incubation in silver nitrate eliminates these artifacts and reduces the commonly observed high background staining. This new method may be valuable for Alzheimer’s disease (AD) researchers and neuropathologists.

## Methods

### Subjects, fixation and paraffin embedding

All subjects or their legal representatives signed an Institutional Review Board-approved informed consent. Slices of brain tissue (one cm thick) are fixed in 10% neutral-buffered formalin for 48 hours at 4 C, dissected into standard tissue cassettes, dehydrated in graded alcohols and xylene and then paraffin-embedded. Paraffin sections (6 μm) are deparaffinized in xylene or xylene substitute and brought through graded alcohols to water.

A comprehensive description of the Civin Laboratory for Neuropathology, Arizona Study of Aging and Neurodegenerative Disorders (AZSAND) and Banner Sun Health Research Institute Brain and Body Donation Program (www.brainandbodydonationprogram.org) is available as a free full-text document^12^ on PubMedCentral.

### Staining method overview

The Bielschowsky silver method was performed with “old” and “new” protocols on 6 µm sections. The old protocol was based on the most commonly-used modification, published by Yamamoto and Hirano in 1996^13^. Up to 24 or 48 slides (the latter when placed in rack back-to-back) are stained in each batch, using plastic 24-slide racks and 200 ml staining containers. All aqueous solutions are made up with reverse-osmosis-purified water; all washes consist of reverse-osmosis purified water.

### Pretreatment with oxidizing agents

The new method differs from the old protocol only in that it includes pretreatment with 0.25% potassium permanganate (Sigma-Aldrich) and 2% oxalic acid (Sigma-Aldrich), trialed at 2, 3 and 5 minutes for each. Slides are washed after these steps in three successive one-minute steps with water.

### Silver nitrate steps

After pretreatment and washing, slides are immersed in 150 ml of a 20% silver nitrate (Sigma-Aldrich) solution for 15 minutes in the dark and then placed in water for five minutes.

The same silver nitrate solution is then titrated by slowly adding 28-30% ammonium hydroxide (Sigma-Aldrich) until the solution clears and no silver precipitate is present. Initially 15 ml of sodium hydroxide are added, then slowly more, drop by drop, until the precipitate that first forms disappears. Finally, 2 more drops are then added to ensure that precipitation has completely disappeared(approximately 22 to 30 ml total). Sections are then placed into this ammonified silver nitrate solution for 10 minutes in the dark. Meanwhile, approximately 10 drops of ammonium hydroxide are added per 100 ml of water to make ammoniacal distilled water. The sections are transferred into the ammoniacal distilled water for 5 minutes prior to adding developer.

### Development and toning steps

The developing agent stock solution is composed of 0.05% formaldehyde (Fisher), 0.003% concentrated nitric acid (Sigma-Aldrich) and 14.27 μM citric acid (Sigma-Aldrich). It is made by adding 4 ml formaldehyde to 25 ml of water, then adding 10 μl concentrated nitric acid and 0.125 g citric acid, then diluting 1:10 in water. This will keep at room temperature for up to two weeks only.

Four ml of developer stock solution are added to the ammonified silver nitrate solution (making up 2.2% of the total volume) from the prior step and the solution quickly stirred; the slides are then placed in this, in the dark, for 5-10 minutes until the section turns a yellowish-tan color or until the required intensity of staining is observed (black fibers with a tan background). The progress may be monitored under the microscope.

Following development, the slides are washed in 3 changes of water for one minute each, and then placed in a 5% sodium thiosulfate (Sigma-Aldrich; 7.5 g in 150 ml water) solution for 30 seconds to remove any unreduced silver from the sections. Slides are then given a final wash in two changes of water for 5 minutes each, dehydrated in alcohols, cleared and coverslipped.

### Comparison of old and new methods

Adjacent sections from blocks that had previously shown plaque-like artifactual staining were stained with both old and new Bielschowsky methods and qualitatively evaluated under the microscope. Additionally, to evaluate whether the new method stained neuritic plaques as well as the old method, adjacent sections with abundant plaques were compared using old and new methods.

To quantitatively compare the ability of the new method to a different gold standard for neuritic plaques, thioflavin S staining^2, 7^, adjacent or semi-adjacent sections of middle frontal gyrus of 30 subjects with varying plaque densities from none to frequent were assessed using the CERAD templates after staining with the new Bielschowsky method and a standard version of the thioflavin S method^14^. Semi-quantitative scores (0-3) for both neuritic plaque and diffuse plaque density were compared using Mann-Whitney U-tests. Spearman rank correlations were calculated, comparing each type of plaque density estimate for both methods, i.e. the new Bielschowsky method versus the thioflavin S method.

## Results

Sections of striatum and cerebellum that had previously and repetitively shown plaque-like artifactual staining with the old method (Figure 1a, c, e) were free of artifact when stained with the new method (Figure 1b, d, f). Pretreatment with 0.25% potassium permanganate and 2% oxalic acid for 2 minutes each was qualitatively judged to give the best results. Longer time trials with pretreatment solutions resulted in damage to tissue integrity. Qualitatively, the new Bielschowsky method (Figure 2b, d) plaques were more readily apparent than with the old method (Figure 2a, c) and background staining, particularly that due to normal myelinated fibers, was reduced. Neurofibrillary tangles were also more readily recognized with the new stain (Figure 3a-d), but this was not quantitatively evaluated.

**Figure 1.**
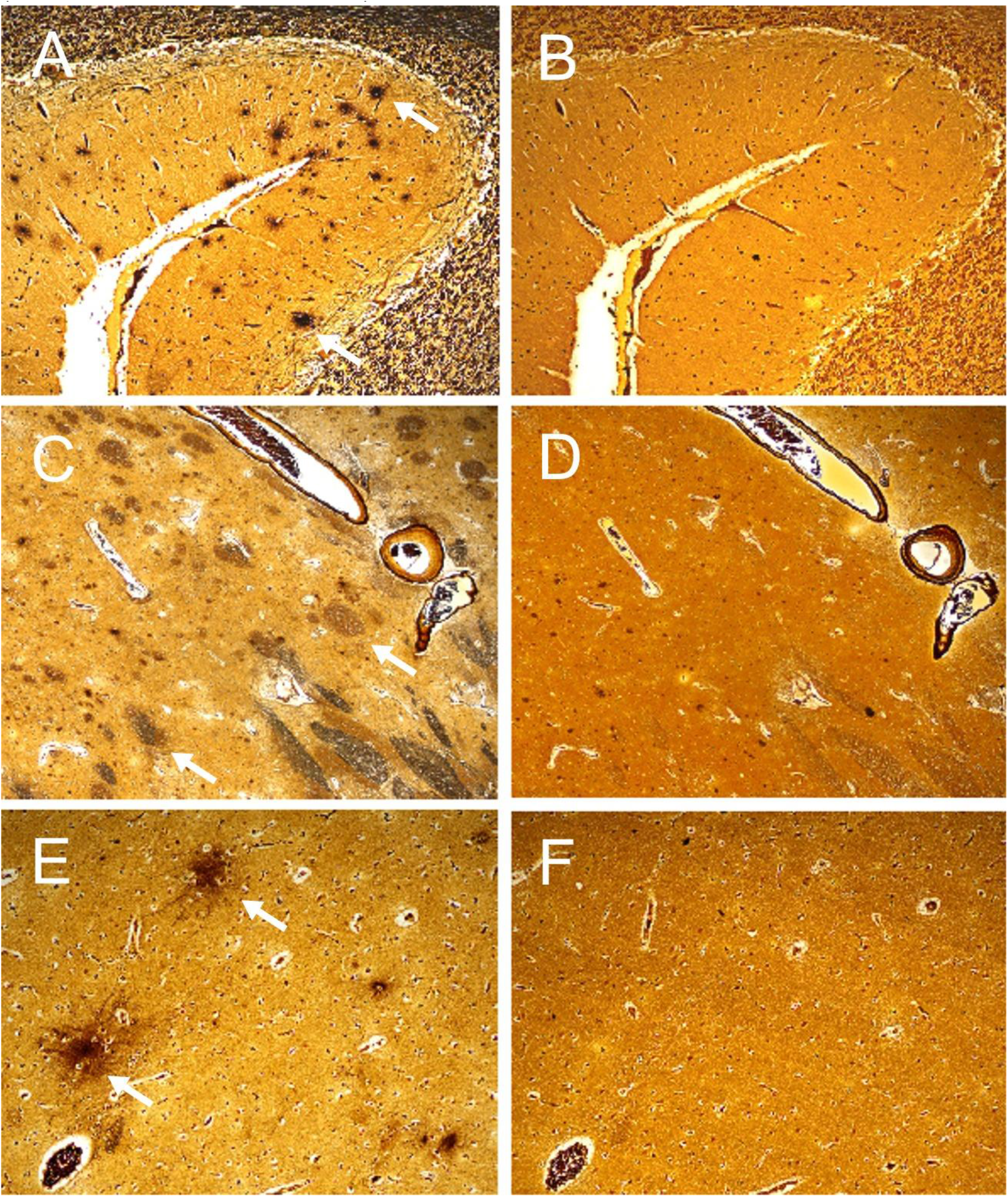
Photomicrographs of adjacent brain sections with few or no true senile plaques, stained with the standard Bielschowsky method of Yamamoto and Hirano^13^ (A, C, E) and the modified Bielschowsky method reported in this manuscript, using pretreatment with potassium permanganate and oxalic acid (B, D, F). Note the artifactual dark spots and darkly-staining normal nerve fiber bundles in the former (arrows point to some examples) and the absence of these with the new stain.

**Figure 2.**
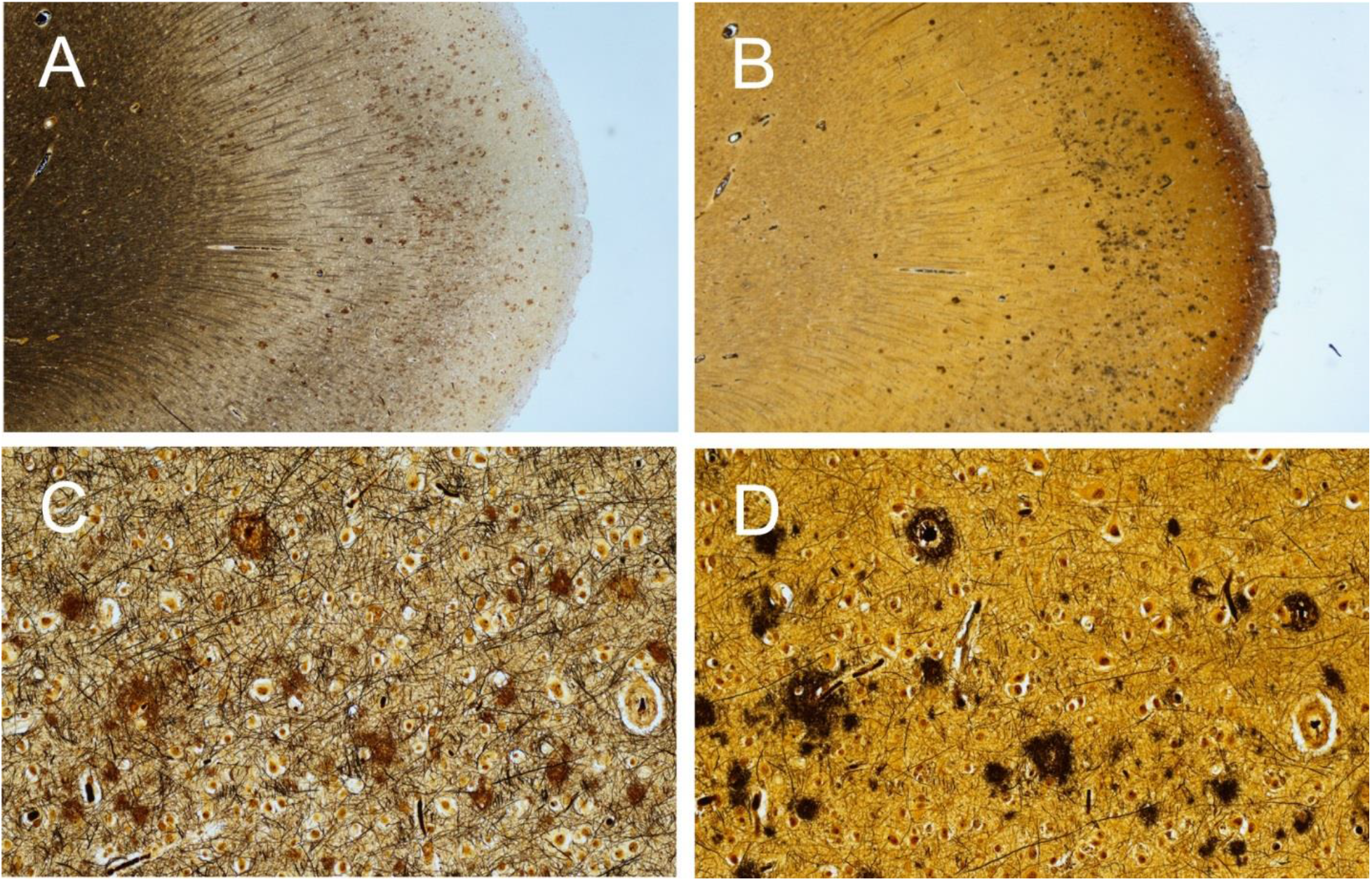
Photomicrographs of adjacent brain sections with abundant senile plaques, stained with the standard Bielschowsky method of Yamamoto and Hirano^13^ (A, C) and the modification reported in this manuscript, using pretreatment with potassium permanganate and oxalic acid (B, D). Note that with the new method, plaques are more readily apparent, and background staining is reduced, at both lower and higher magnifications.

**Figure 3.**
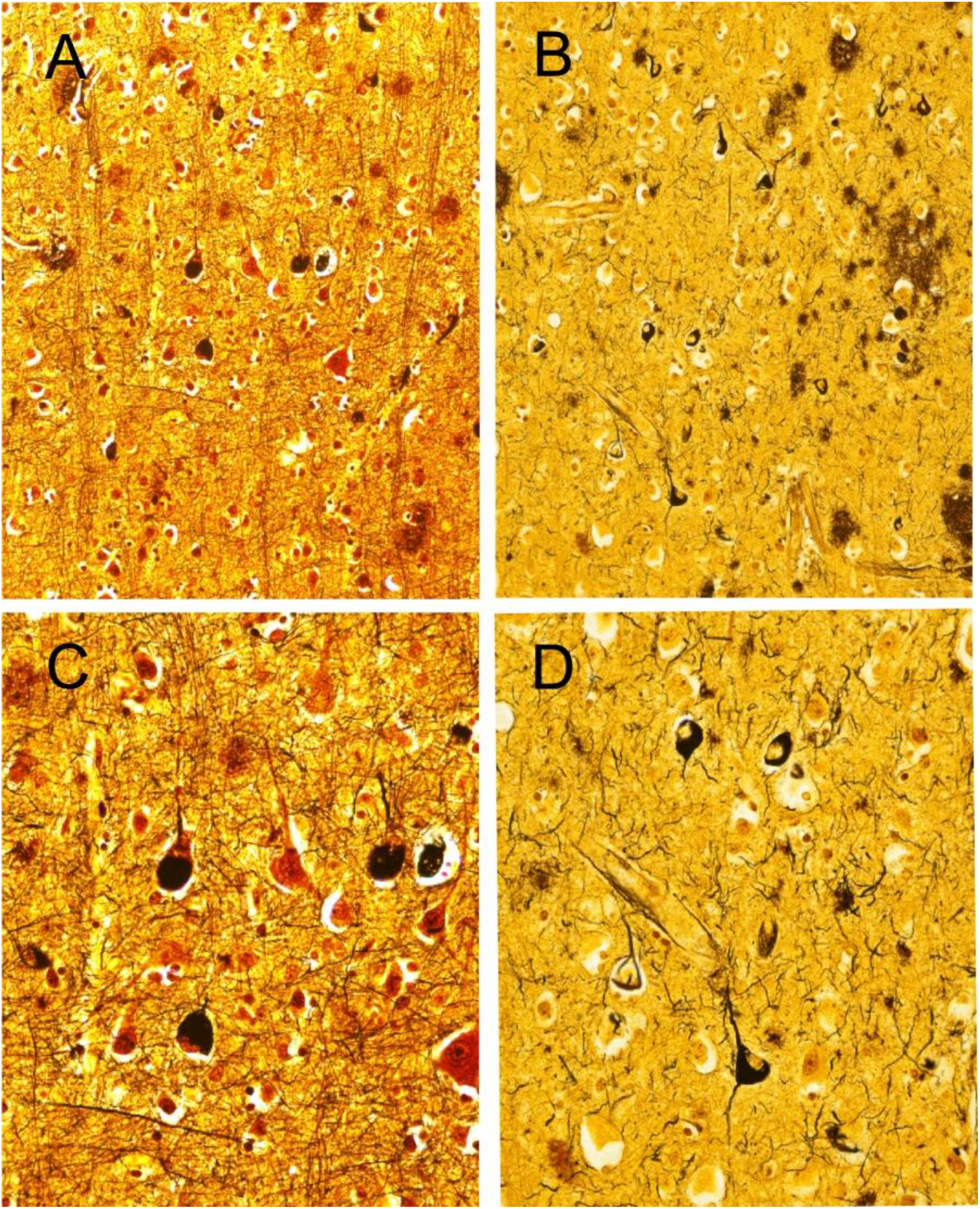
Photomicrographs of semi-adjacent brain sections stained with a standard Bielschowsky method based on that of Yamamoto and Hirano^13^ (A, C) and the modification reported in this manuscript, using pretreatment with potassium permanganate and oxalic acid (B, D). Note that with the new method, neurofibrillary tangles and neuropil threads are more readily apparent, and background staining is reduced, at both lower and higher magnifications.

Quantitatively, plaque density estimates for either neuritic or diffuse type did not significantly differ between the new Bielschowsky method and the thioflavin S method (Table 1). Spearman correlations were greater than 0.9 and highly significant (p < 0.0001) for both neuritic and diffuse plaques stained with the two methods.

**Table 1.** Comparison of semi-quantitative estimates of neuritic and diffuse plaque densities using the new Bielschowsky method and a standard thioflavin S method. Means and standard deviations are given. The two methods did not differ significantly, demonstrating that the new stain identifies both plaque types as well as a gold standard method, the thioflavin S stain.

| **Neuritic Plaques**<br>**New Bielschowsky** | **Neuritic Plaques**<br>**Thioflavin S** | **Diffuse Plaques**<br>**New Bielschowsky** | **Diffuse Plaques**<br>**Thioflavin S** |
| --- | --- | --- | --- |
| 1.67 (1.21) | 1.73 (1.26) | 1.93 (1.28) | 2.0 (1.36) |
| Mann-Whitney U-test $p = 0.77$ | | Mann-Whitney U-test $p = 0.70$ | |
| Spearman $\rho = 0.94$; $p < 0.0001$ | | Spearman $\rho = 0.92$; $p < 0.0001$ | |

## Discussion

Pretreatment of paraffin sections with potassium permanganate and oxalic acid eliminated troublesome artifactual staining with the Bielschowsky method. Such artifactual staining is common in sections with little or no true AD pathology. It is possible that the artifactual staining results from the localized presence of reducing agents in the tissue that precipitate elemental silver. We reasoned that pretreatment with oxidizing agents would strip electrons from these sites, preventing them from reducing silver ions in subsequent steps. The oxidizing agents were chosen by analogy with their usage in the Gallyas silver method^1, 15-17^. We are not aware of any previous Bielschowsky modifications that have employed oxidizing agent pretreatment for this purpose. Lhotka et al (1953) experimented on the ability of five oxidizing agents to differentiate normal from degenerating nerve fibers^18^. They employed varying concentrations of incubation times with potassium permanganate, chromic acid, periodic acid, lead tetra-acetate and sodium bismuthate, but not oxalic acid.

This modification of the Bielschowsky stain allows the confident determination of the absence of plaques in a tissue section, by eliminating artifactual staining that mimics the appearance of plaques. Due to reduction of the background staining from normal nerve fibers, both plaques and tangles are more readily appreciated. As compared with the gold standard thioflavin S method for neuritic plaques, this new Bielschowsky method does not produce differing estimates of either neuritic or diffuse plaque densities, assuring that it may be substituted for the commonly-used Bielschowsky method of Yamamoto and Hirano^13^.

## Acknowledgements

The Civin Laboratory for Neuropathology, Arizona Study of Aging and Neurodegenerative Disorders and Brain and Body Donation Program have been supported by the National Institute on Aging (P30 AG19610 Arizona Alzheimer’s Disease Core Center), the National Institute of Neurological Disorders and Stroke (U24 NS072026 National Brain and Tissue Resource for Parkinson’s Disease and Related Disorders), the Arizona Department of Health Services (contract 211002, Arizona Alzheimer’s Research Center), the Arizona Biomedical Research Commission (contracts 4001, 0011, 05-901 and 1001 to the Arizona Parkinson’s Disease Consortium) and the Michael J. Fox Foundation for Parkinson’s Research.

